# A Facile Method for Simultaneous Visualization of Wet Cells and Nanostructured Biomaterials in SEM using Ionic Liquids

**DOI:** 10.1101/2020.08.05.238246

**Authors:** Bryan E.J. Lee, Liza-Anastasia DiCecco, Hourieh Exir, Arnaud Weck, Kyla Sask, Kathryn Grandfield

**Affiliations:** School of Biomedical Engineering, McMaster University, Hamilton, Canada; Department of Materials Science and Engineering, McMaster University, Hamilton, Canada; Department of Physics, University of Ottawa, Ottawa, Canada; Centre for Research in Photonics at the University of Ottawa, Ottawa, Canada; Department of Mechanical Engineering, University of Ottawa, Ottawa, Canada

**Author notes:** Corresponding Author: Kathryn Grandfield, Department of Materials Science and Engineering, McMaster University, 1280 Main Street West, Hamilton, ON, L8S 4L7, Canada.

**Keywords:** ionic liquid, wet cell SEM, cell adhesion, nanostructures, titanium, implant

## Abstract

This work presents a successful methodology to image mammalian cells adhered to nanostructured biomaterials using scanning electron microscopy (SEM) operating in low-vacuum mode following ionic liquid treatment. Human osteoblast-like Saos-2 cells were treated with a room-temperature ionic liquid, 1-Ethyl-3-methylimidazolium tetrafluoroborate, and subsequently imaged on titanium utilizing SEM. Titanium substrates were modified to create laser-induced periodic surface structures (LIPSS) for visualizing at the sub-micron scale. Using a combination of fluorescence-based cell metabolism along with light microscopy and SEM image analysis, the shape and location of irradiated cells were confirmed to be unchanged after multiple irradiation sessions while the viability of minimally irradiated cells was unaltered. The wet imaging conditions combined with a rapid facile protocol using ionic liquid allows this technique to fulfill a niche in examining cellular behavior on biomaterials with sub-micron surface features. The demonstrated method to track observed cell adhesion to sub-micron surface features with SEM has great implications for the understanding of cell migration on nanostructured surfaces as well as on the exploration of simpler SEM preparation methods for cellular imaging.

## Introduction

The performance of any biomaterial is contingent on its interactions between its surface and cells.^[1]^ Identifying the shape and morphology of these cells is often indicative of how the biomaterial may perform in the future. However, as biomaterials innovation moves towards surfaces with micro- to nanoscale surface topography, these features can currently only be observed using electron microscopy methods. This visualization is particularly important for the biomedical device industry which frequently uses sub-micron or nanoscale features to improve cellular outcomes such as anodized nanotubes,^[2]^ laser-induced periodic surface structures (LIPSS),^[3]^ or nanopatterned grooves.^[4]^ While the advent of super-resolution techniques has enabled optical techniques to achieve nanoscale resolution of cells via the creative use of fluorescent tools, this has encroached upon the territory once owned by electron microscopy.^[5]^ Unfortunately, these optical techniques only apply for components that can be fluorescently tagged. This means that the visualization of all other objects is limited to the diffraction of light or to images at the micron-scale. As such, to allow for simultaneous imaging of the interface between cells and sub-micron substrates, electron microscopy remains the best choice. Nanostructures are used in dynamic biomedical applications ranging from implant materials, biosensors, and tissue engineering to drug delivery.^[6–9]^ The proliferation and adhesion of cells in each of these applications and how they change over time is often tied to the overall success of the biomaterial.^[10]^ Therefore, a method to image cells on these surfaces would facilitate better biomaterial design through the visualization of the direct cell to micro/nano-scale feature interactions.

Imaging of wet or living biological samples that are viable using electron microscopy is a great challenge that has yet to be overcome. The mechanism by which the electron microscope works is, in of itself, incredibly detrimental to the viability of biological specimens. In the case of transmission or scanning transmission electron microscope (TEM/STEM), it has been considered almost impossible to image living biological cells although attempts have been made by encapsulating fixed cells in between silicon nitride windows.^[11,12]^ There have been attempts to reduce the most harmful effects of scanning electron microscopy (SEM) by reducing the accelerating voltage, operating under low-vacuum, or utilizing environmental microscopes (ESEM).^[13,14]^ However, these techniques still struggle to image cells without causing irreversible damage. Chemical fixation, dehydrating, and staining are other commonly employed methods to image cells but these techniques come with the drawback of altering the morphology of cells, proteins, and other relevant molecules resulting in the imaging of a sample which has been altered from its living state.^[15]^ These dried samples lack the fluidity of cells in their native environment and thus are not completely representative of the system.

Within the last decade, attempts have been made using novel room-temperature ionic liquid (RTIL) treatments to image biological cells using SEM. Specifically, RTILs are described to have unique properties such as minimal vapour pressure as well as high conductivity.^[16]^ This combination of properties allows RTILs to be used instead of conventional metal coatings for non-conductive samples and in particular RTILs can provide a conductive layer for biological samples while wet or in liquid form. In microbial imaging applications, RTIL treatment of unfixed hydrated samples has been found to provide comparable imaging quality and contrast to traditional dehydration preparation methods paired with staining or conductive coating.^[17]^ Moreover, RTIL treatment can mitigate artifacts in biological samples such as wrinkling, shrinkage, and cracking, and it may provide a closer morphological representation of the natural hydrated microbe state.^[17]^ Researchers have utilized RTILs purely as an alternative to metal sputter coating,^[17–20]^ while others have used RTIL to image living bacterial cells and red blood cells with SEM.^[21,22]^ While these studies have shown the capacity to image cells, they have often been limited to single endpoint studies on non-complex substrates with simple organisms such as bacteria. Additionally, literature currently available that utilize RTILs in techniques have focused on cells that were already fixed or dehydrated, with limited works exploring live samples. Of importance for biological cells and medical devices are the interactions that occur at the cell-substrate interface.

This work utilizes a unique facile RTIL treatment using 1-Ethyl-3-methylimidazolium tetrafluoroborate diluted with cell culture media to simultaneously image both mammalian cells and the LIPSS titanium sub-micron surface features they were adhered to using SEM. Cell SEM imaging was subsequently evaluated using biochemical assays and light microscopy to confirm cellular viability following both RTIL treatment and SEM imaging. The minimal vapour pressure of the RTIL used combined with low-vacuum SEM imaging was able to facilitate cellular observation under liquid conditions, which is a more representative model of the native cellular environment. This paper demonstrates RTIL treatment can facilitate imaging of cells in liquid conditions to study textured surface interactions using SEM.

## Materials and Methods

### Titanium Sample Preparation

Commercially pure Grade 2 titanium was cut using a lathe and blade setup to produce disks with a diameter of 15 mm and a thickness of 1.25 mm. Titanium disks were polished using a four-step procedure in which the titanium was exposed, in order, to silicon-carbon sandpaper, 9 μm and 3 μm diamond polishing suspensions, and for final polishing a colloidal silica suspension (OPS) mixed with 10% H_2_O_2_. Laser modified disks were prepared using a Yb:KGW femtosecond laser as outlined in previous work.^[3]^ All disks were ultrasonically cleaned for 15 minutes in both ethanol and acetone. Titanium disks were scratched using a dremel tool to create distinct features to allow for tracking of cell migration.

### Cell Culturing and Metabolism

Saos-2, osteosarcoma, cells (ATCC ®, HTB-85) were grown in McCoy’s 5A modified media (Life Technologies Inc.) with 15% fetal bovine serum (Life Technologies Inc.) and 1% penicillin/streptomycin (Sigma-Aldrich). Cells were incubated at 37oC with 5% CO_2_. Cells were seeded on the titanium samples while placed in 12-well plates and allowed to adhere for 1 day before imaging. For longer-term viability experiments, the media was exchanged every day following cell metabolism experiments.

### Scanning Electron Microscope Imaging using RTIL

The RTIL process is highlighted schematically in Figure 1A. The culture media was removed from wells containing samples and was then replaced with a 5% ionic liquid solution (in McCoy’s 5A modified media). The RTIL used was 1-Ethyl-3-methylimidazolium tetrafluoroborate (Sigma-Aldrich). The ionic liquid solution was left in the well for 5 minutes before being aspirated, after which samples were removed from the well plate and dried gently on both sides through blotting. Samples were then mounted onto 6-inch diameter SEM stubs using carbon tape and then placed in the SEM for observation.

**Figure 1:**
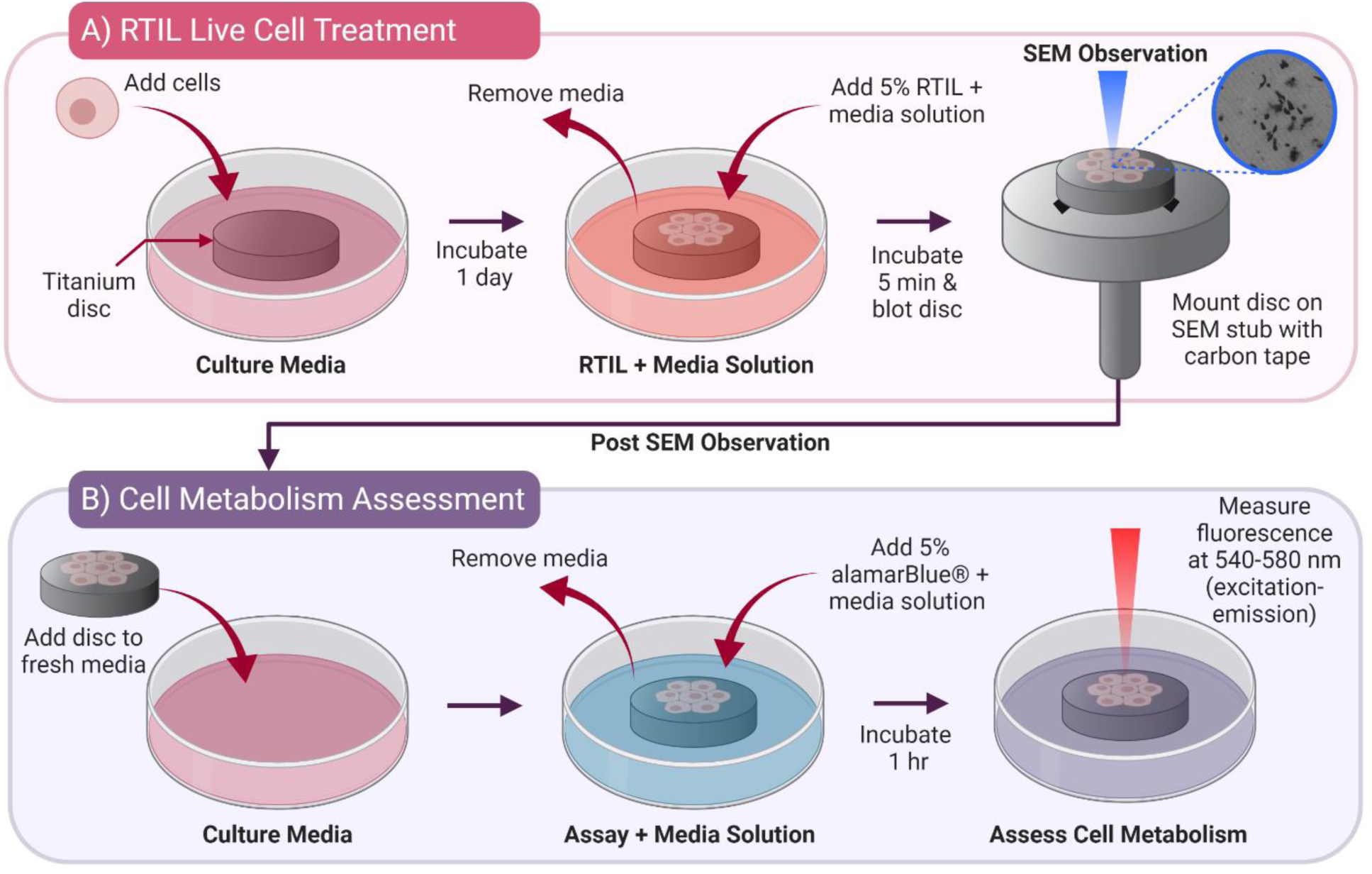
Schematic demonstrating the method for RTIL treatment and cell assessment. A) Preparation of samples for imaging. B) Evaluation of cell metabolism following imaging in SEM. Figure created with BioRender.com.

A TESCAN VP SEM (Tescan, Czech Republic) was operated under two different conditions. Primarily, the SEM was operated in low-vacuum mode using a backscattered electron detector with an accelerating voltage of 10kV. For specific experiments, SEM imaging was also operated under traditional high-vacuum conditions utilizing secondary electron acquisition. Samples were imaged for no longer than 30 minutes at a time, no more than three times a day. In between experiments samples were kept in an incubator at 37°C and 5% CO_2_. All measurements from SEM images were obtained using ImageJ (National Institute of Health).

### Cell Metabolism after RTIL and SEM

Cell metabolism was measured using an alamarBlue® (Life Technologies Inc.) assay. As described schematically in Figure 1B, following imaging under the SEM, cells were rinsed with PBS before a 5% alamarBlue solution (in McCoy’s 5A media) was added to each well. The samples were incubated in the dark for 1 hour at 37°C before obtaining fluorescent readings using an Infinite ® M1000 (Tecan, Männedorf, Switzerland) at 540-580 nm (excitation-emission). Blank readings were subtracted from each data point and data was normalized by cells grown in untreated wells. In non-endpoint studies, following fluorescent reading, the alamarBlue® solution was removed and replaced with media, and samples were placed back in the incubator at 37°C with 5% CO_2_. These samples would subsequently be treated with RTIL and imaged using SEM at a later date.

### Cell Viability after RTIL, SEM and Cell Metabolism

To determine the long-term viability of the cells following both RTIL treatment and SEM imaging, cells were detached from titanium samples using trypsin in 0.25% EDTA (Sigma–Aldrich) as per ATCC guidelines and media was added to deactivate trypsin after detachment was observed. Cells were then replated onto 12-well tissue culture plates and allowed to grow on the plates. Some samples, before imaging, were stained with a Nile Red stain (ThermoFisher Scientific). A solution of 10 mL PBS and 10 μL Nile Red was added to the re-plated cells and left to incubate for 10 minutes before being removed. Images of the cells were taken using an Olympus IX51 Inverted microscope (Olympus, United States of America).

## Results and Discussion

Initial treatment of the cells with the RTIL, growing in tissue culture plates, was determined to overall be non-toxic to cells, which were able to grow and proliferate following RTIL treatment (Figure 2). There were no statistically significant differences (p < 0.05) in cell metabolism (Fig 2 A) between cells that were RTIL treated. This is qualitatively and visibly observed in the stained cells that were incubated following initial RTIL treatment (Fig 2 B/C). While some RTILs have shown various degrees of toxicity towards cells in short term exposure,^[16]^ these results demonstrate that 1-Ethyl-3-methylimidazolium tetrafluoroborate in the quantities and approach used in this work is non-toxic to Saos-2 cells.

**Figure 2:**
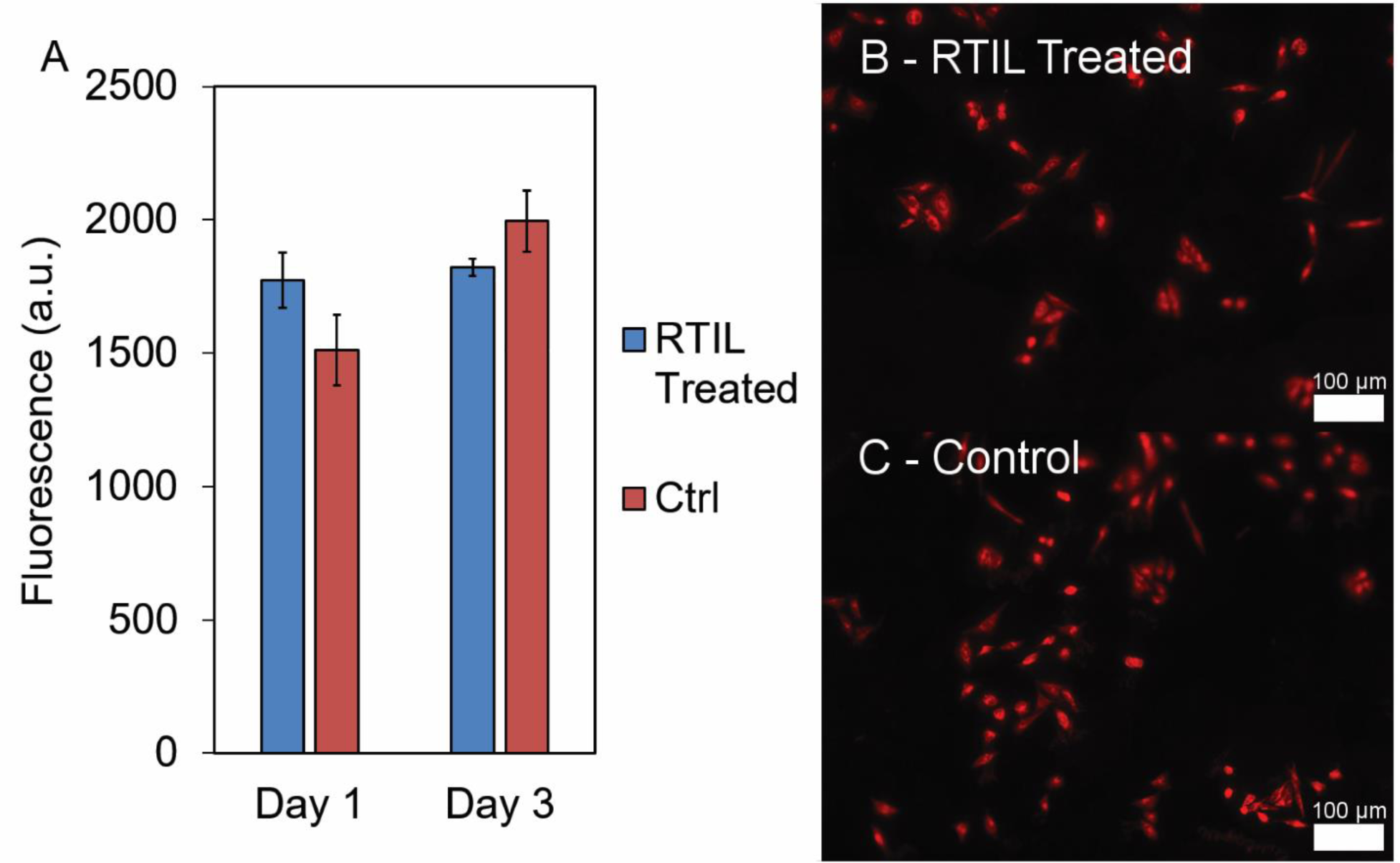
Cell metabolism data for cells plated on 12 well plates that were either exposed to RTIL for 5 minutes or unaltered. There was no statistically significant (p < 0.05) difference in cell metabolism of treated and non-treated cells (A). Cells were stained with Nile Red after 3 days and show comparable morphology in RTIL compared to control treatments.

The ability to visualize cells adhered to opaque materials with sub-micron features is a specific niche that can only be solved with electron microscopy. Cells adhered to titanium substrates with and without sub-micron features are observed using SEM in Figure 3. Numerous cells can be seen with an elongated morphology which is indicative of their viability when placed in the microscope. Additionally, the RTIL treatment also provides some contrast, even in backscattered mode, as the RTIL becomes trapped in the sub-micron features which allow for the generation of some topographical contrast. This pooling has been noted in other work with RTILs and has been speculated to be a result of variations in viscosity of the solution.^[19,23]^ Fixation or dehydration protocols paired with coating or staining could also be used as conventional approaches to imaging Saos-2 cells on the LIPSS titanium surface, however, these preparation techniques do not allow for hydrated sample observation and are lengthy in process, ranging from hours to days of sample preparation compared to the RTIL treatment used which takes only minutes. While low magnification images reveal confluent or highly concentrated areas of cells (Fig 3A/B), the higher magnification images (Fig 3C/D) show that cells may be interacting with the sub-micron features. While low-vacuum backscattered imaging was utilized for all imaging in this work, there are pros and cons to utilizing other imaging modes such as secondary electron imaging in higher vacuum conditions or high-vacuum ESEM. Studies have shown that radiation damage is greater in the low-vacuum mode, but general cell viability is noted to be worse in high-vacuum methods.^[24]^ Moreover, the difference in vapour pressures in the high-vacuum ESEM is believed to contribute to a large driving force for mass transfer of water out of cells.^[25,26]^ Conversely, low-vacuum imaging has been shown to increase the concentration of water molecules which may increase the rate of specimen degradation observed.^[25,26]^ Therefore, in this study, we have used a combination of low-vacuum along with the described RTIL treatment to enable cell imaging with minimal damage to the cells.

**Figure 3:**
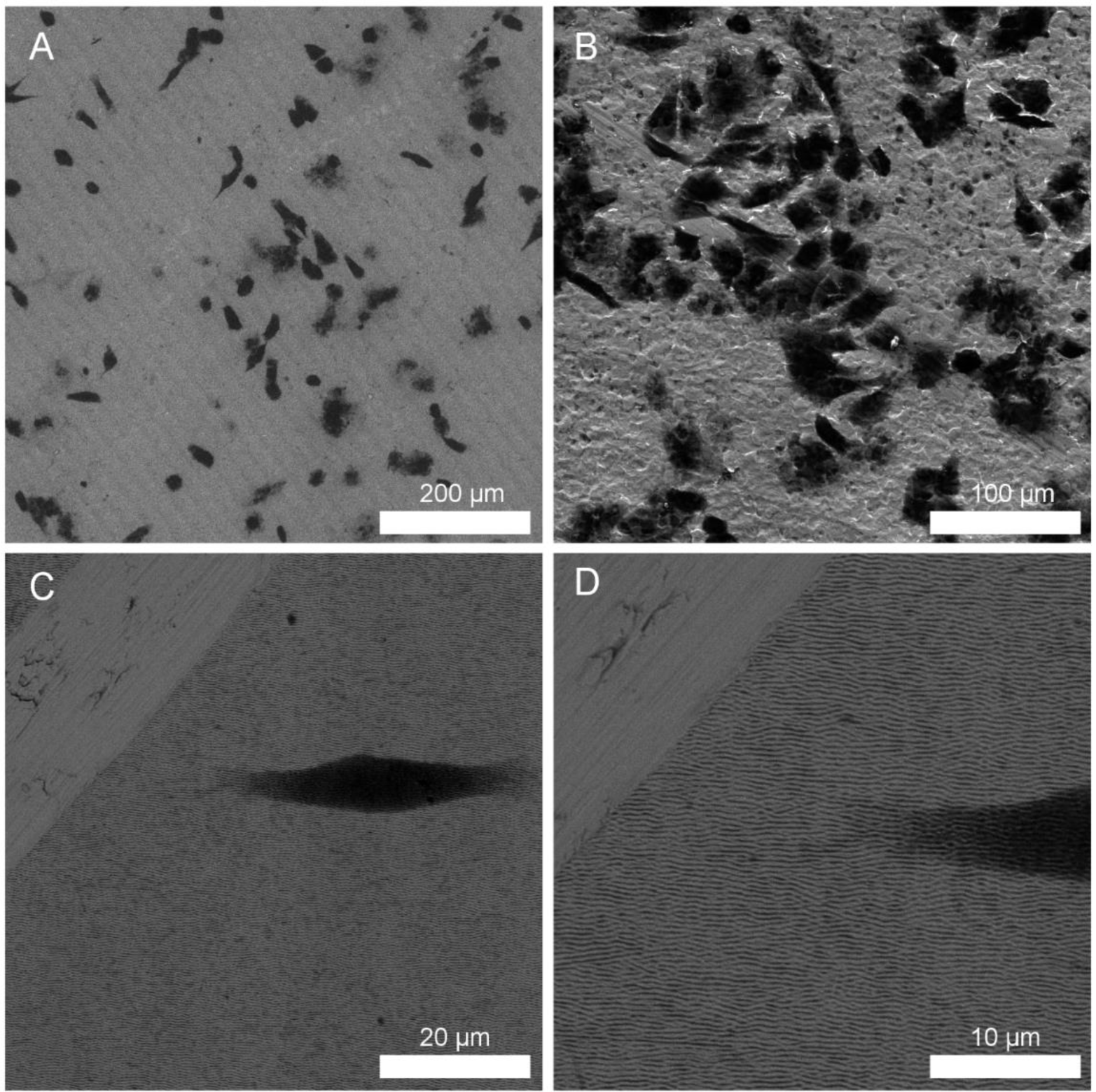
Cells adhered to titanium substrates after ionic liquid treatment for 5 minutes. Cells can be observed across the entire substrate (A/B) and in detail with respect to sub-micron features (C/D).

The adhesion and migration of cells along the surfaces of biomaterials is essential to understand cell-material interactions. Here, cells were seeded on titanium substrates with laser-induced periodic surface structures (LIPSS)^[3]^ and visualized after 1 (Figure 4A) and 3 (Figure 4B) days. From previous work, it was determined that these laser modified surfaces have periodicities between 300 and 620 nm with an average roughness of 145 nm.^[3]^ Tracking the position of the same cells over multiple days was possible because of the large, macroscale scratches that were applied with a dremel tool. In theory, any consistent macroscale feature could be utilized to aid in the tracking of cells, be it naturally occurring in the material or artificially added. The cells remained intact and viable from 1 to 3 days as per corresponding cell metabolism readings (Table 1). However, the differences in cell position may be too small to be considered meaningful in this case and imaging artifacts may contribute. Subsequent viability experiments indicate that cell death occurred to cells exposed directly to the electron beam following RTIL treatment and placement in the SEM. Notably, the cells appear identical in shape and location in micrographs up to 3 days following initial RTIL treatment and SEM irradiation at 1 day and then a subsequent repeat at 3 days. This demonstrates that while the cells may no longer be viable, their location and shape remain intact, and thus this quick treatment can be used multiple times on the same sample within this set time frame, with preservation of imaging quality. Research involving RTIL treatment on fixed cells on the other hand has shown a downfall in that it is not permanent either. While cells are preserved and imaged well up until five days post-RTIL treatment, after seven days they may start to experience charging in SEM and subsequently imaging quality will decrease.^[20]^

**Table 1:** Cell to cell and scratch to cell measurements taken in ImageJ demonstrating how the cells have migrated across the laser modified surface over time. Cell metabolism readings indicate that cells are healthy after multiple RTIL treatments and multiple SEM sessions. (1) – scratch to close cell, (2) – scratch to far cell, (3) – cell to cell correspond to measurements indicated in Figure 4.

| Substrate | Time Point | (1) - Scratch to Close Cell ($\mu\text{m}$) | (2) - Scratch to Far Cell ($\mu\text{m}$) | (3) - Cell to Cell ($\mu\text{m}$) | Cell Metabolism (a.u.) |
| --- | --- | --- | --- | --- | --- |
| LIPSS Titanium | Day 1 | 3.6 | 42.3 | 22.9 | 736 |
|  | Day 3 | 4.8 | 43.4 | 26.1 | 742 |
|  | Change | 1.2 | 1.1 | 3.2 | N/A |
| Unmodified Titanium | Day 1 | 98.5 | 177.1 | 33.9 | 740 |
|  | Day 3 | 98.5 | 177.1 | 34.1 | 686 |
|  | Change | 0 | 0 | 0.2 | N/A |

**Figure 4:**
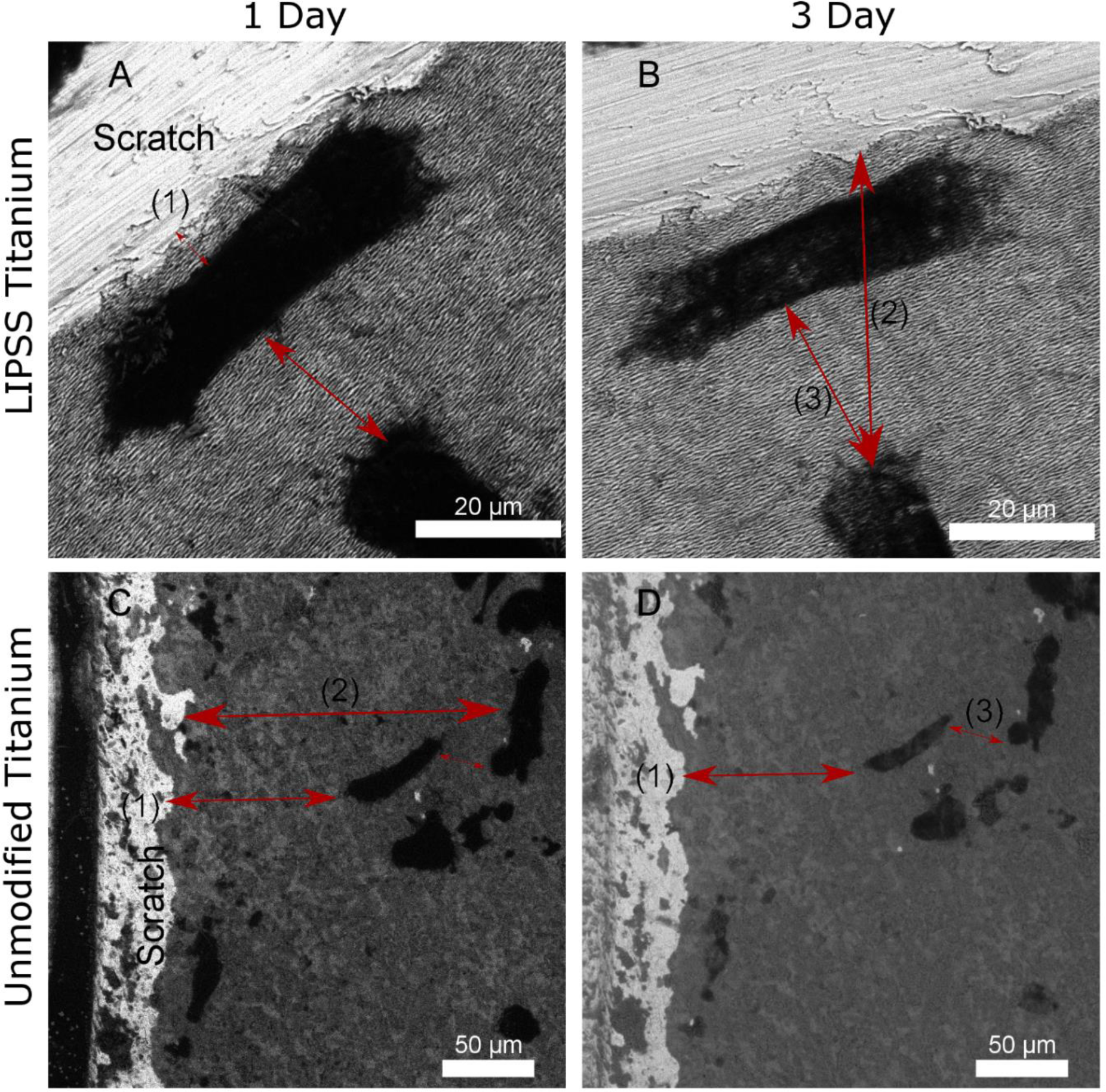
Demonstration of cells visualized on laser ablated (A/B) and unmodified (C/D) titanium observed using SEM with ionic liquid treatment alongside correlated fluorescent cell metabolism readings. The difference in position of the cells in (A/C) compared to (B/D) show how the cells are unchanged in shape or location following multiple SEM irradiations sessions and RTIL treatments. (1) – scratch to close cell, (2) – scratch to far cell, (3) – cell to cell.

A major concern of utilizing electron microscopy to image biological materials is the risk of irreversible damage to the cells. As such, following RTIL treatment and SEM imaging, cells were detached from their titanium substrates and re-plated onto tissue-culture plates to confirm that the cells maintained their viability and phenotype. Figure 5 shows cells imaged with an inverted light microscope 1 (Figure 5 A) and 3 (Figure 5 B/C) days following re-plating. These images show adhered cells with an elongated morphology which demonstrates that the cells grow and proliferate under typical cell culture conditions following repeated RTIL treatment and SEM imaging. Figure 5C shows cells that were stained by Nile Red, a dye that stains lipids.^[27,28]^ The cell membrane and numerous organelles have lipid bilayers which would have been ruptured if damage from the electron beam or the vacuum was fatal.^[11]^ The cells are well stained with the membranes visibly intact suggesting that damage from this procedure was minimal or negligible. This confirms that while directly irradiated cells were rendered unviable, cells in the periphery or those that were only briefly irradiated were not altered by RTIL treatment or by SEM imaging. This is confirmed by the cell metabolism readings (Table 1) observed after 1 and 3 days which show that the overall viability of the surrounding cells is unaltered. Previous work on cells adhered to LIPSS titanium identified that cells tended to be adhered perpendicular to the alignment of the LIPSS^[3]^ but this study using RTIL treatment did not observe the same mechanisms. The cells observed in previous work^[3]^ were prepared using a lengthy sample preparation process involving fixation, dehydration, staining, and coating and thus it is possible that only certain cells remained properly attached to the LIPSS following the sample preparation steps and that the morphology of cells was altered through conventional preparation means. Thus, the RTIL treatment presents an advantage over more traditional biological SEM sample preparation as cells are still in liquid conditions and are not fixed or dehydrated. Further improvements to biological imaging in the SEM could be achieved via software implemented rastering protocols, as shown in beam irradiation studies of epoxy resin.^[29]^

**Figure 5:**
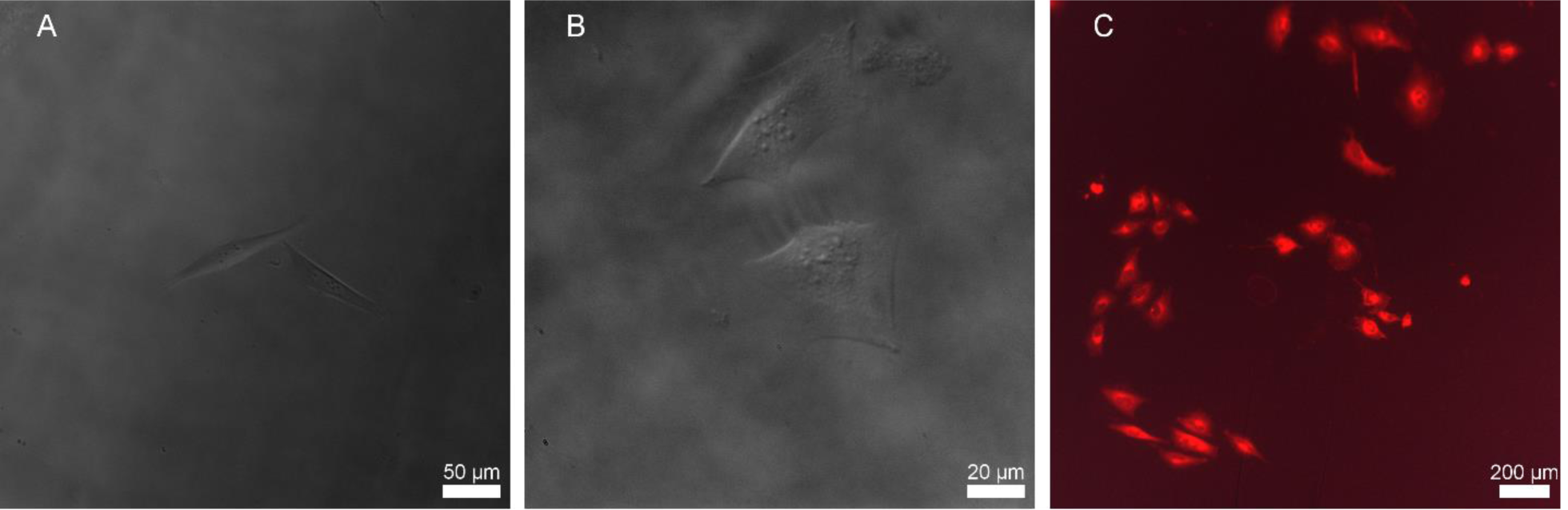
Demonstration of the viability of cells following RTIL treatment and imaging, in low-vacuum mode, in the SEM. Cells appear elongated which is indicative of cellular viability following treatments. Cells were detached from titanium substrates with trypsin and re-plated into empty 12 well plates. Cells were imaged 1 (A) and 3 (B/C) days after re-plating. (C) shows cells that were stained by Nile Red, a lipid stain.

The compatibility of biomaterials is highly associated with cellular adhesion and migration along their surfaces. This work demonstrates the power of using these treatments to evaluate cell adhesion to sub-micron features on modified materials such as those modified with LIPSS. Cell adhesion and migration along surfaces are examples of a phenomenon that requires equal attention to both the cells and the surface itself, and, herein, we have demonstrated a method to simultaneously image both with the nanometer resolution afforded by SEM. While this work has demonstrated the capacity for imaging of wet cells on metallic substrates, the RTIL treatment developed for mammalian cells can be expanded more broadly to other cell-surface applications that require wet conditions, such as to understand cell interactions with three-dimensional scaffolds, to monitor the effect of pharmaceuticals on cells or for various other materials and surface modification techniques. Examination of the specific interactions of cells with micro to nanostructured surfaces will contribute to information essential to the design of future biomaterials, drug delivery platforms, and tissue-engineered scaffolds.

## Conclusions

This work presents a new RTIL treatment to facilitate the imaging of wet mammalian cells in the SEM. Herein, we showed that a solution of RTIL (1-Ethyl-3-methylimidazolium tetrafluoroborate) and cell media was non-cytoxic to mammalian Saos-2 cells on LIPSS titanium. The RTIL treatment provided sufficient contrast and charge disappation for SEM imaging of cells on nanostructured surfaces, thus avoiding timely fixation and dehydration protocols. After irradiation in the SEM, cells were replated and displayed their regular phenotype. This technique can therefore fill an important niche providing a facile approach for imaging both cells and their nanostructured interfacing surfaces with the sub-diffraction limit resolution afforded by SEM.

## Acknowledgments

We would like to acknowledge the support of the Natural Sciences and Engineering Research Council of Canada (NSERC) and the Discovery Grant Program (RGPIN 2014-06053). Microscopy was carried out at McMaster Faculty of Health Sciences Electron Microscopy facility. *In vitro* studies were performed at the Biointerfaces Institute at McMaster University.

## Conflicts of Interest

The authors declare no conflict of interest.

## Data Availability

Data is available upon request to the corresponding author.

## Author Contributions

**Conceptualization:** Bryan E.J. Lee, Kathryn Granfield

**Methodology:** Bryan E.J. Lee, Kathryn Granfield

**Data curation:** Bryan E.J. Lee, Hourieh Exir

**Formal analysis:** Bryan E.J. Lee, Hourieh Exir, Kathryn Granfield

**Funding acquisitions and supervision:** Arnaud Weck, Kyla Sask, Kathryn Granfield

**Writing:** Bryan E.J. Lee, Liza-Anastasia DiCecco, Kyla Sask, Kathryn Granfield

**Review and editing:** Bryan E.J. Lee, Liza-Anastasia DiCecco, Hourieh Exir, Arnaud Weck, Kyla Sask, Kathryn Granfield

## Notes

### Competing Interest Statement

The authors have declared no competing interest.

